# Evaluation of a multiplexed tiling PCR scheme for whole-genome amplification of hepatitis B virus using Oxford Nanopore sequencing

**DOI:** 10.64898/2026.03.28.714721

**Authors:** Jon Bråte, Eugenia Giovanna Grande, Benedikte Nevjen Pedersen, Torstein Gjølgali Frengen, Kathrine Stene-Johansen

## Abstract

Here we evaluated the performance of a previously published tiling PCR primer scheme by Ringlander et al. (2022) for whole-genome amplification of Hepatitis B virus (HBV) in combination with Oxford Nanopore sequencing. The primer set originally developed for Ion Torrent sequencing was adapted by removing platform-specific adapters and tested using clinical serum or plasma samples submitted for routine HBV genotyping and resistance testing. Two multiplexing strategies were compared: a single PCR pool containing all primers and a two-pool strategy with non-overlapping amplicons. Sequencing reads were processed using a Nanopore analysis pipeline, and genome coverage and amplicon performance were compared across samples spanning a wide Ct range and representing HBV genotypes A–E.

Across all samples, the median genome coverage was approximately 50%, although recovery varied widely, ranging from complete failure to nearly full genomes. Combining all primers into a single PCR reaction, or separating overlapping amplicons into different reactions, had little overall impact on genome recovery, and no consistent differences between the two pooling strategies were observed. In contrast, amplification efficiency differed markedly between individual amplicons. Amplicons 1–5 generally produced higher sequencing depth, whereas amplicons 6–10 frequently showed low coverage and contributed to incomplete genome recovery. Genome coverage was strongly associated with Ct values, with higher coverage observed in samples with lower Ct values, while coverage was broadly similar across genotypes.

These results demonstrate that the Ringlander et al. primer scheme can be adapted for multiplex PCR and Nanopore sequencing of HBV, but uneven amplicon performance limits consistent full-genome recovery and highlights the need for further optimization of HBV tiling PCR designs.

## Introduction

Hepatitis B virus (HBV) remains a major global health concern, causing more than a million deaths annually, primarily due to liver cirrhosis and hepatocellular carcinoma (WHO 2024). Although widespread vaccination programs have greatly reduced incidence in many countries, chronic HBV infection continues to represent a significant public health challenge, particularly among populations from high-prevalence regions. Chronic infection may be treated with antiviral therapy, yet long-term management is dependent on continued genomic monitoring of circulating strains to detect the emergence of novel mutations and drug resistance (Kim et al. 2014).

Historically, genomic sequencing of HBV has relied on the amplification of partial genomic regions, most commonly the polymerase and surface genes, followed by Sanger sequencing (e.g., Kim et al. 2014; Ma et al. 2011; Naito et al. 2001). However, partial genome sequencing fails to capture variation across the entire genome and can miss clinically relevant mutations in other regions, such as the core and precore regions of the HBV e antigen. Whole-genome sequencing (WGS) provides a more comprehensive view of viral diversity and enables more accurate genotype or sub-genotype assignment, while also facilitating detection of co-infections, recombination events, and resistance mutations (Rajoriya et al. 2017). In addition to supporting clinical characterization of individual infections, whole-genome data can also contribute to molecular surveillance by enabling detailed monitoring of circulating HBV lineages and emerging variants.

The high specificity and sensitivity of PCR make it suitable for targeted enrichment of HBV DNA from clinical samples. However, because of the high genomic diversity among HBV strains, reliable primer design that can capture the full diversity is difficult (Chen et al. 2023; Rajoriya et al. 2017). Tiling PCR, where multiple overlapping amplicons spanning the genome are combined, has therefore become a widely used strategy for viral whole-genome sequencing. Each amplicon covers a portion of the genome, and together they provide redundancy: if one amplicon fails, the others can still recover much of the genome. This approach gained prominence during the Zika virus epidemic (Quick et al. 2017) and was refined in the ARTIC Network protocol during the COVID-19 pandemic.

Despite the widespread use of tiling PCR for viral whole-genome sequencing, its application to HBV remains limited. Only a small number of studies have proposed multi-amplicon schemes tailored to the HBV genome. Hebeler-Barbosa et al. (2020) designed three overlapping amplicons optimized for Illumina sequencing, whereas Ringlander et al. (2022) developed a set of ten overlapping amplicons of approximately 400-500 bp that covered the entire HBV genome. More recently, Lumley et al. (2025) designed a scheme consisting of six longer amplicons (∼600-700 bp), but with large primer redundancy at each site to better capture the HBV genetic diversity.

Although the primer scheme designed by Ringlander et al. (2022) demonstrated successful and highly sensitive amplification across several HBV genotypes (A, B, C, and D), it was only sequenced using the Ion Torrent platform. Furthermore, the primers contained Ion Torrent-specific adapter sequences, limiting their compatibility with other sequencing technologies. Long-read sequencing technologies such as Oxford Nanopore offer substantial advantages for viral genomics, including the recovery of complete amplicons in single reads, which simplifies genome assembly and improves haplotype reconstruction. Lumley et al. (2025) successfully applied Nanopore sequencing to sequence full-length amplicons from their tiling PCR scheme. Although the Ringlander et al. (2022) amplicons are somewhat shorter, adapting this primer set for Nanopore sequencing would still be advantageous. However, it remains uncertain whether these primers perform efficiently without the Ion Torrent adapters, or whether they can be multiplexed at all.

In tiling PCR workflows, two key multiplexing strategies are commonly used. One approach pools all primers into a single PCR reaction, as done in Integrated DNA Technologies (IDT) protocols. This has the advantage that only a single PCR reaction is required per sample. But risk the amplification of primarily very short fragments from the overlapping amplicons. The other approach, as used in the ARTIC Network method protocols for example, divides primers into two pools of non-overlapping amplicons. While this requires two PCR reactions per sample and increases both the time of the workflow and the risk of sample mixup, it avoids the primer competition and formation of short overlapping fragments.

In this study, we therefore evaluated the performance of the Ringlander et al. (2022) primer set for whole-genome amplification of HBV using Oxford Nanopore sequencing. Specifically, we tested whether the primers can (1) be used without Ion Torrent adapters, (2) be combined into multiplex PCR reactions using either a single pool or two non-overlapping pools, and (3) successfully amplify HBV genomes across multiple genotypes, including genotype E.

## Materials and Methods

### Samples and DNA extraction

Serum or plasma samples were submitted to the Norwegian Institute of Public Health (NIPH) mainly for routine HBV genotyping and resistance testing as part of the patient management. In selected cases, samples were also analyzed for detection of vaccine escape mutations or for characterization of rare, conflicting, or atypical diagnostic markers.

The samples analyzed in this study had been previously sequenced for a small (∼800bp) region where the P and S-genes overlap using PCR and Sanger sequencing. The resulting sequence was genotyped using Geno2Pheno (Beggel et al. 2009). Ct values were determined by quantitative PCR following the protocol described by (Pettersson et al. 2019). We selected in total 84 different samples representing genotypes A (30 samples), B (19 samples), C (16 samples), D (37 samples – two of these were part of a EQA panel from QCMD), and E (18 samples) that were analyzed in this study (see Supplementary Table S1 for details). 36 of these samples were run in parallel to test the impact of different primer pooling approaches (described below), which makes for a total of 120 different analyses. DNA was extracted from 200 µl plasma/serum using the Qiagen QIAamp DNA Mini Kit (Qiagen, Netherlands) on the QIACube (Qiagen, Netherlands) automated system following the manufacturers protocol or using the Abbott sp2000 (Abbott, IL, US). Extracted DNA was eluted in 100 µL elution buffer and stored at −70°C prior to PCR amplification.

### Primer set and pooling strategies

The primers designed by Ringlander et al. (2022) consists of ten pairs producing overlapping amplicons across the HBV genome (∼400 bp each). The primers were originally described with sequencing adapters for Ion Torrent. We removed these sequences to only retain the HBV-binding binding regions (Figure 1 and Table 1). Primer binding sites were estimated using the Geneious Prime v2024.0.7 software against the well annotated and widely used NC_003977.2 HBV genotype D reference sequence (Table 1).

**Figure 1:**
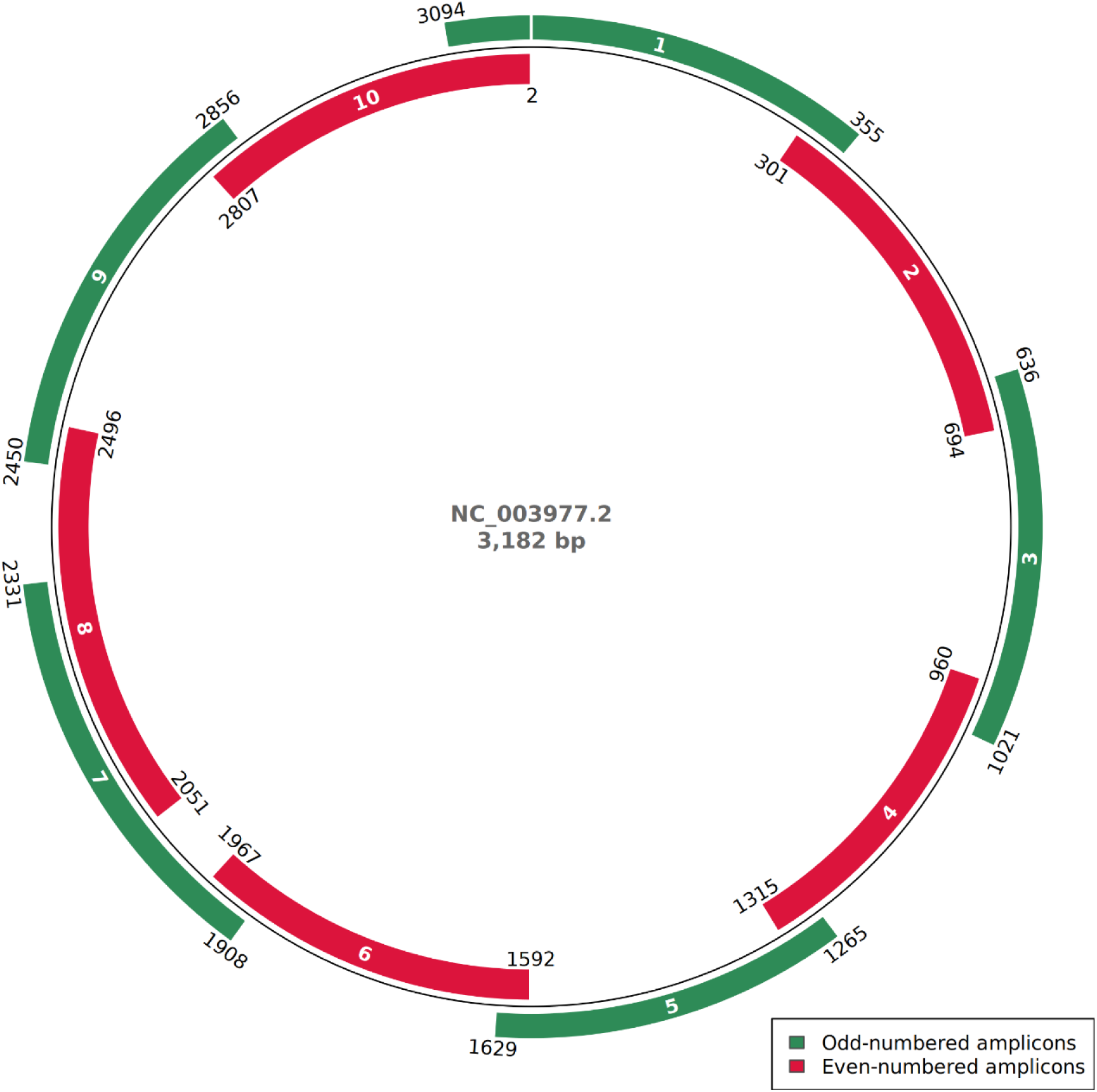
The regions amplified by the modified Ringlander et al. (2022) primers as estimated by mapping the primers to the HBV genotype D reference sequence NC_003977.2. Amplicons are numbered 1-10 as in the original publication.

**Table 1:**
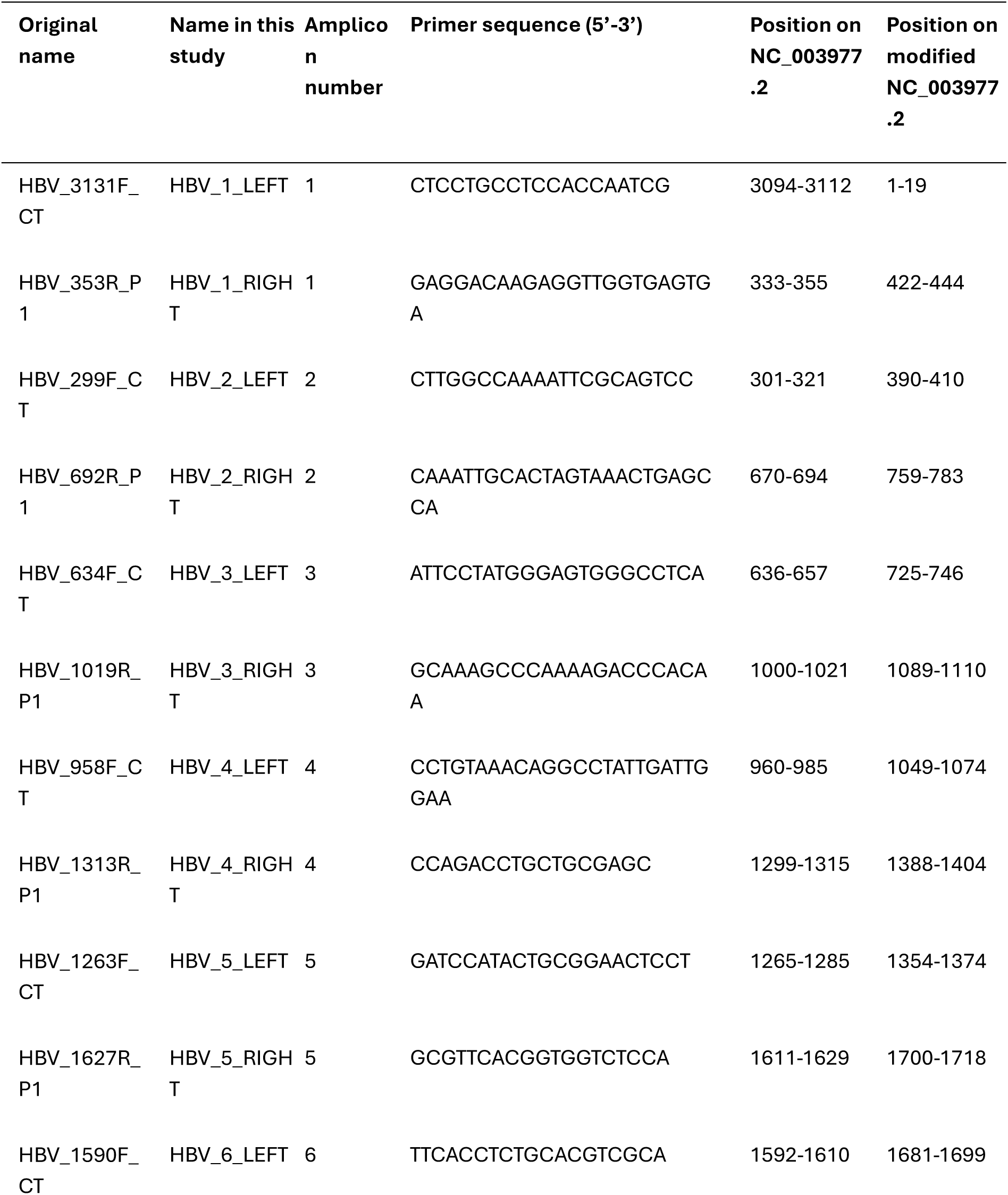
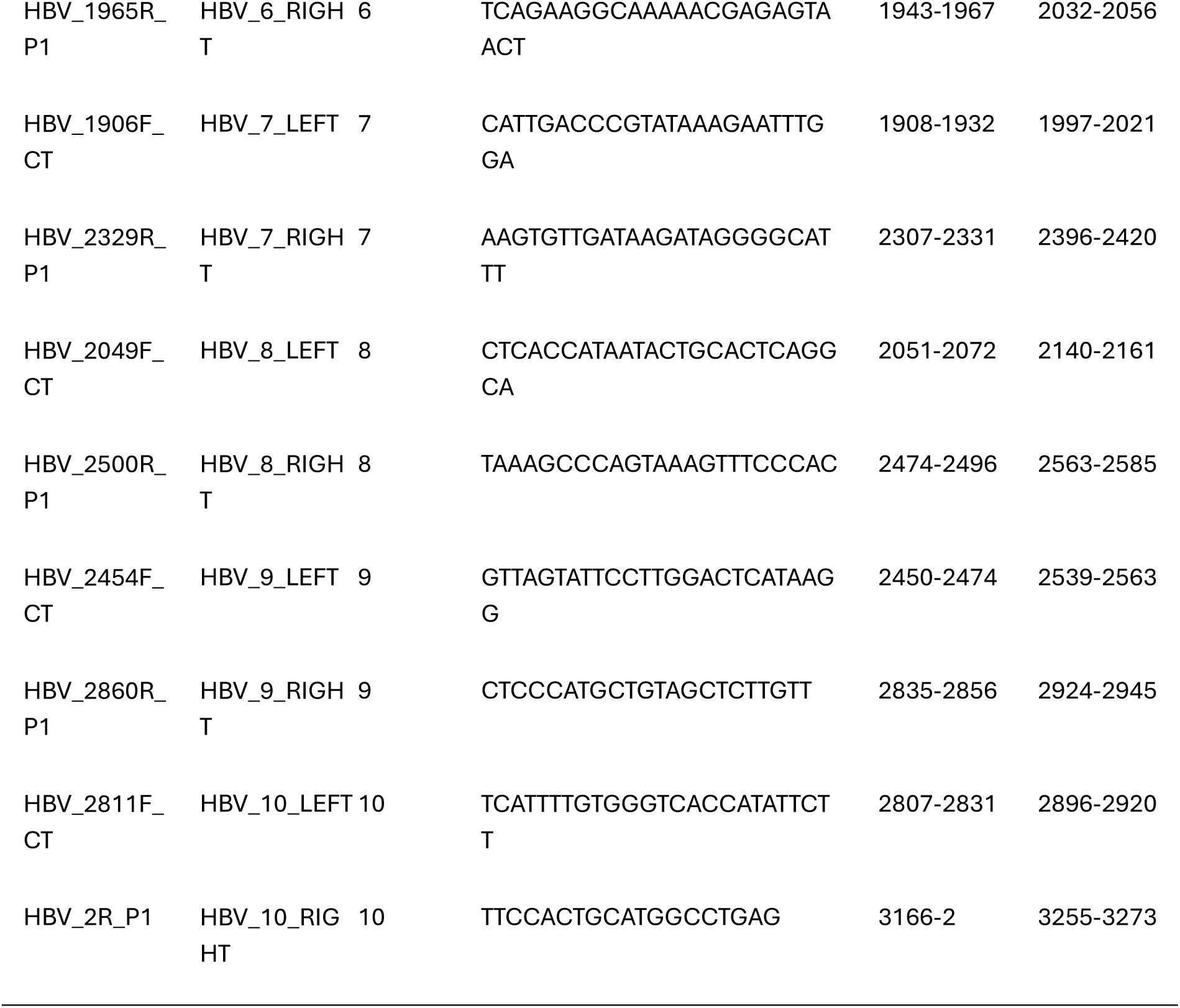
Primer sequences and binding sites. “Original name” is the names used by Ringlander et al. 2022. “Name in this study” is the primer names we used in our analyses. “Amplicon number” is the number given to the amplicons by Ringlander et al. 2022 and the number used in our analyses. “Primer sequence (5’-3’) is the sequence of the primers we used in our study (i.e., IonTorrent adapters removed). “Position on NC_003977.2” is the estimated primer binding site on the original reference sequence. “Position on modified NC_003977.2” is the estimated primer binding site on the modified reference sequence (see description in Materials and Methods).

36 samples (Supplementary Table S1) were tested using two different primer pooling strategies:

#### One primer pool

all 20 primers were combined into a single pool (one PCR reaction per sample) with a total primer concentration of 20 µM.

#### Two primer pools

the primers were separated into two pools with non-overlapping amplicons (two PCR reactions per sample). Pool 1 consisted of primers for odd-numbered amplicons and pool 2 of primers for the even-numbered amplicons (Table 1). Each pool had a total primer concentration of 10 µM.

### PCR amplification

Amplification was performed using Platinum SuperFi II DNA Polymerase (Thermo Fisher Scientific, MA, US) with the following cycling conditions: initial denaturation at 98°C for 30 s; 35 cycles of 98°C for 10 s, 60°C for 30 s, and 72°C for 3 min; and a final extension at 72°C for 5 min. Amplicons were visualized on TapeStation D5000 (Agilent, CA, US) and quantified by Qubit dsDNA HS assay (Thermo Fisher Scientific).

### Nanopore library preparation and sequencing

Amplicons were prepared for sequencing using the Oxford Nanopore Technologies (ONT) Rapid Barcoding Kit (SQK-RBK114-96) and sequenced on GridION Mk1 using R10.4.1 flow cells (ONT, UK). In total, four different sequencing runs were performed. Two for the samples amplified with a single primer pool (named HBV9 and HBV10) and two for the samples amplified with two primer pools (HBV11 and HBV12; see Supplementary Table S1).

### Read processing, mapping and generation of consensus sequences

Basecalling and demultiplexing were performed using the MinKNOW software v24.02.16 with Dorado v7.3.11 (ONT, UK) and the high accuracy 400 bps model. Reads were processed using version 1.8.5 of the ARTIC pipeline (https://github.com/artic-network/fieldbioinformatics) which can handle libraries produced by the Nanopore Rapid kit. Reads were first demultiplexed and quality filtered using *guppyplex* (part of the ARTIC pipeline) with only reads between 100-4000 bp retained. Next, reference mapping and consensus sequence generation was done using *minion* as part of the ARTIC pipeline. In brief, reads were mapped using *minimap2* (Li 2018) against the modified version of reference NC_003977.2 (see below), primer sequences were trimmed after mapping, and consensus sequences were created using *Clair3* (Zheng et al. 2022) with the model r1041_e82_400bps_hac_v500 and a minimum read depth of 20 reads to call a consensus nucleotide. The *minion* pipeline also estimates the normalized abundance of each amplicon which were used to compare the performance of each amplicon. For the *minion* pipeline we used two different primer bed-files, one for each primer pooling strategy (see Supplementary File 1). Consensus sequences were genotyped using HBV-GLUE online version (Singer et al. 2018).

*minimap2* takes as input a linear reference sequence. However, because the HBV genome is circular, linearizing the reference sequence means that reads spanning the origo (i.e., originating from amplicon 1 and 10) map at both ends of the reference and hence must be fragmented. To avoid this, we modified the reference sequence by simply duplicating the region of overlap between amplicon 1 and 10. We also set the 1^st^ nucleotide of the linearized sequence to be the start of amplicon 1, and the last nucleotide to be the end of amplicon 10 (see Supplementary File 1). The estimated primer binding sites on the modified sequence are shown in Table 1.

### HBV read classification

The sequenced reads were classified as either HBV or non-HBV using Kraken2 v2.1.6 (Wood et al. 2019). The HBV Kraken2 database was created by downloading all available HBV sequences with a minimum length of 2500 bp (117 182 sequences) from the HBV-GLUE website (https://hbv-glue.cvr.gla.ac.uk/#/home. Data downloaded September 3. 2025. The file is available as Supplementary File 2) and building a custom Kraken2 database from these.

### Human read removal

Prior to submission of the read files to the European Nucleotide Archive, all FASTQ files were screened for human-derived reads using an alignment-based approach. Reads were aligned to the T2T-CHM13v2.0 complete human reference genome assembly using minimap2 (v2.24+) with the Oxford Nanopore preset (*map-ont*). The T2T-CHM13v2.0 assembly was selected over GRCh38 following Forbes et al. (2025), who demonstrated that the telomere-to-telomere assembly achieves substantially higher human read removal sensitivity due to its inclusion of previously unresolved genomic regions. Reads that did not align to the human reference (SAM flag-f 4) were retained and written to a new gzip-compressed FASTQ file using SAMtools v1.22.1. The number of input reads, human reads removed, and output reads were logged for each sample (Supplementary Table S1).

### Primer mismatch analysis

Selected reference sequences representing each HBV genotype were obtained from the study by (McNaughton et al. 2020). To identify the theoretical primer-binding region in each reference sequence, a sliding-window matching approach was implemented in R v4.3.3. For each primer, each reference sequence was scanned using a window of size equal to the primer length, advancing one nucleotide at a time. At each position, the number of mismatches between the primer sequence and the corresponding reference subsequence was calculated by direct sequence comparison. Both the forward strand and the reverse complement of each reference sequence were evaluated, and the position with the lowest number of mismatches was retained as the best local match. A maximum of 10 mismatches were allowed. For the best local match, the number of mismatches against the entire primer was counted, in addition to the last 5 nucleotides to represent 3’-end mismatches. This approach does not take into account potential insertions or deletions.

### Data availability

The supplementary data files for this publication are available under the FigShare project named “HBV tiling PCR”. This project contains the following items:

Supplementary_File_1: Bed-files and modified reference sequence used for the ARTIC pipeline (doi: 10.6084/m9.figshare.31684666).

Supplementary_File_2: HBV genome sequences used to build a Kraken2 database for sequence classification (doi: 10.6084/m9.figshare.31577149).

The Nanopore sequence data (fastq files) with human reads removed (described above) will be deposited to the Europoean Nucleotide Archive under the Project PRJEB110761 upon publication of this manuscript.

## Results

### Sequencing output

The sequencing, read classification and mapping results for all samples (negative controls excluded) are summarized in Table 2. Note that these statistics are generated from all 120 analyses, including 36 samples that were sequenced in duplicate (see Materials and Methods). Sequencing output varied substantially, with raw read counts ranging from 575 to 415,509. Similarly, both the proportion of reads classified as HBV and the mapping rate to the reference genome showed considerable variation across samples. The majority of reads passed *Guppyplex* quality of length filtering.

**Table 2:** Sequencing, classification and mapping statistics. “Raw reads” are the fastq_pass reads produced by the MinKnow software. “Classified reads” are the reads classified by Kraken2 against an HBV-specific database. “Classification rate (%)” is the fraction of raw reads classified by Kraken2. “Guppyplex filtered” is the number of reads retained after Guppyplex. “Mapped reads” is the number of Guppyplex filtered reads mapping to the reference using minimap2. “Mapping rate (%)” shows the fraction of Guppyplex filtered reads that mapped.

| Processing stage | Mean ± SD | Median | Min | Max |
| --- | --- | --- | --- | --- |
| Raw reads | 29 137 ± 47 156 | 16 756 | 575 | 415 509 |
| Classified reads | 18 959 ± 23 972 | 10 146 | 35 | 168 650 |
| Classification rate (%) | 66.9 ± 31.3 | 78.5 | 0.5 | 98.3 |
| Guppyplex filtered | 28 201 ± 46 412 | 16 145 | 550 | 407 357 |
| Mapped reads | 16 746 ± 21 938 | 8687 | 31 | 164 375 |
| Mapping rate (%) | 62.6 ± 28.3 | 73.8 | 0.4 | 99.7 |

### Effect of differential primer pooling

We first tested the impact on whole-genome amplification of pooling all primers into a single amplification reaction versus separating them into two reactions containing non-overlapping amplicons. 36 samples were analyzed using both primer pooling strategies. Overall, genome coverage showed a positive correlation between the two approaches (Figure 2A), with a Pearson correlation coefficient of R = 0.68 (p < 0.001), indicating a moderate-to-strong association.

**Figure 2:**
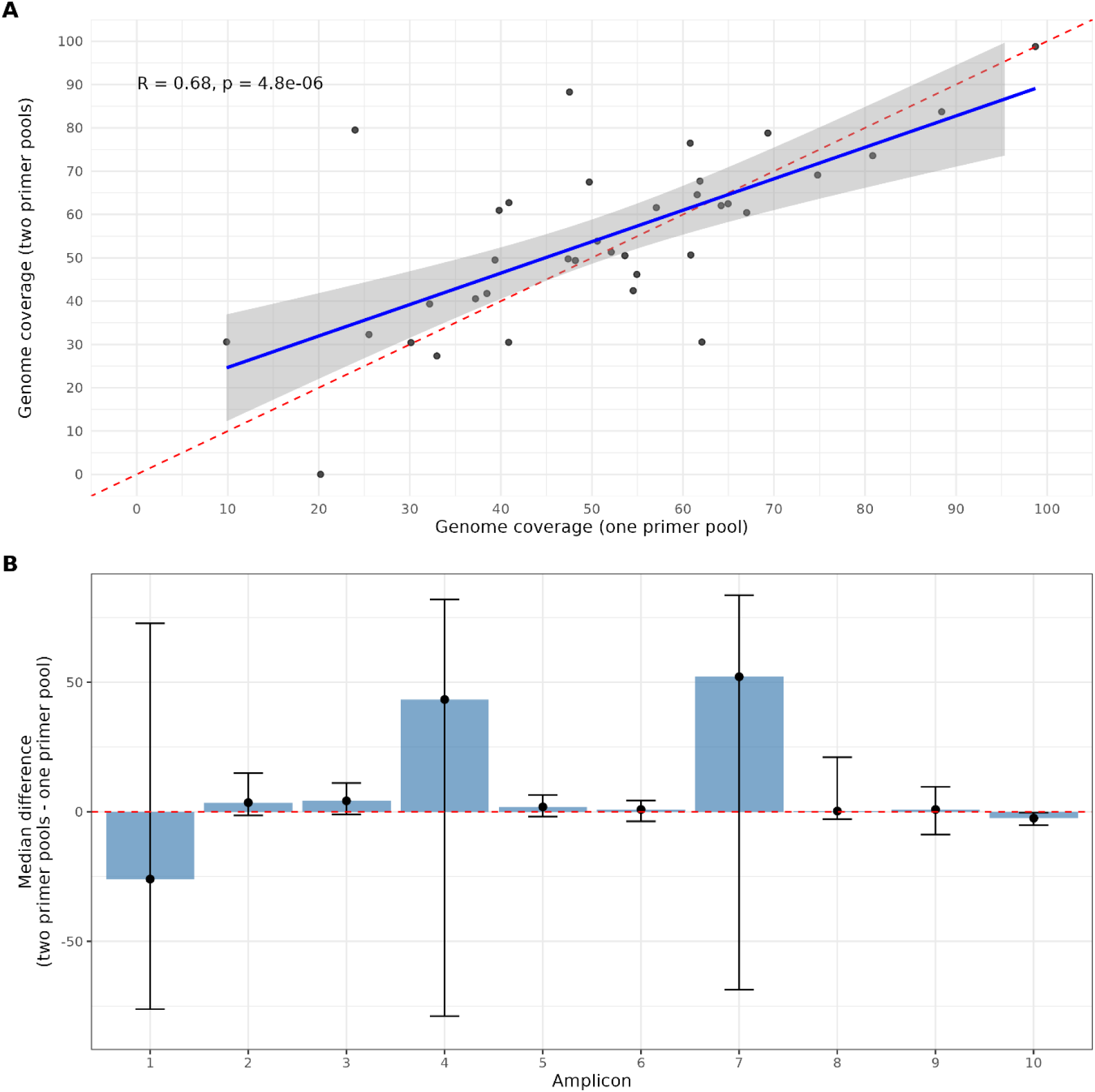
A) Pairwise comparison of genome coverage in percentage between samples analyzed with both one and two primer pools. Red dashed line indicates the perfect correlation. The blue line is the linear regression fitted to the data with the 95% confidence intervals for the regression shown as shaded area. *R* shows the Pearson correlation with the probability of this correlation being due to chance indicated by *p*. B) Comparison of normalized amplicon depths between samples analyzed with both primer pooling strategies. The blue bars represent the median difference (marked also by a black dot) in amplicon depths between the two primer pooling strategies. The range of the differences is represented by the error bars (25^th^ to 75^th^ percentile).

### Genome recovery

Next, we analyzed genome coverage across all samples regardless of primer pooling strategy (Figure 3). Samples analyzed multiple times are plotted independently. Genome coverage varied considerably, ranging from 0% (completely failed) to 99% (full coverage), with a median of 50% and an interquartile range of 32–63%. While approximately half of the samples achieved >50% genome coverage, only two samples exceeded 90% coverage, and 25% of samples had coverage below 32%. Coverage was similar across genotypes, with median values ranging from 42% (genotype A) to 62% (genotype C). The substantial variability within each genotype exceeded the differences between genotypes.

**Figure 3:**
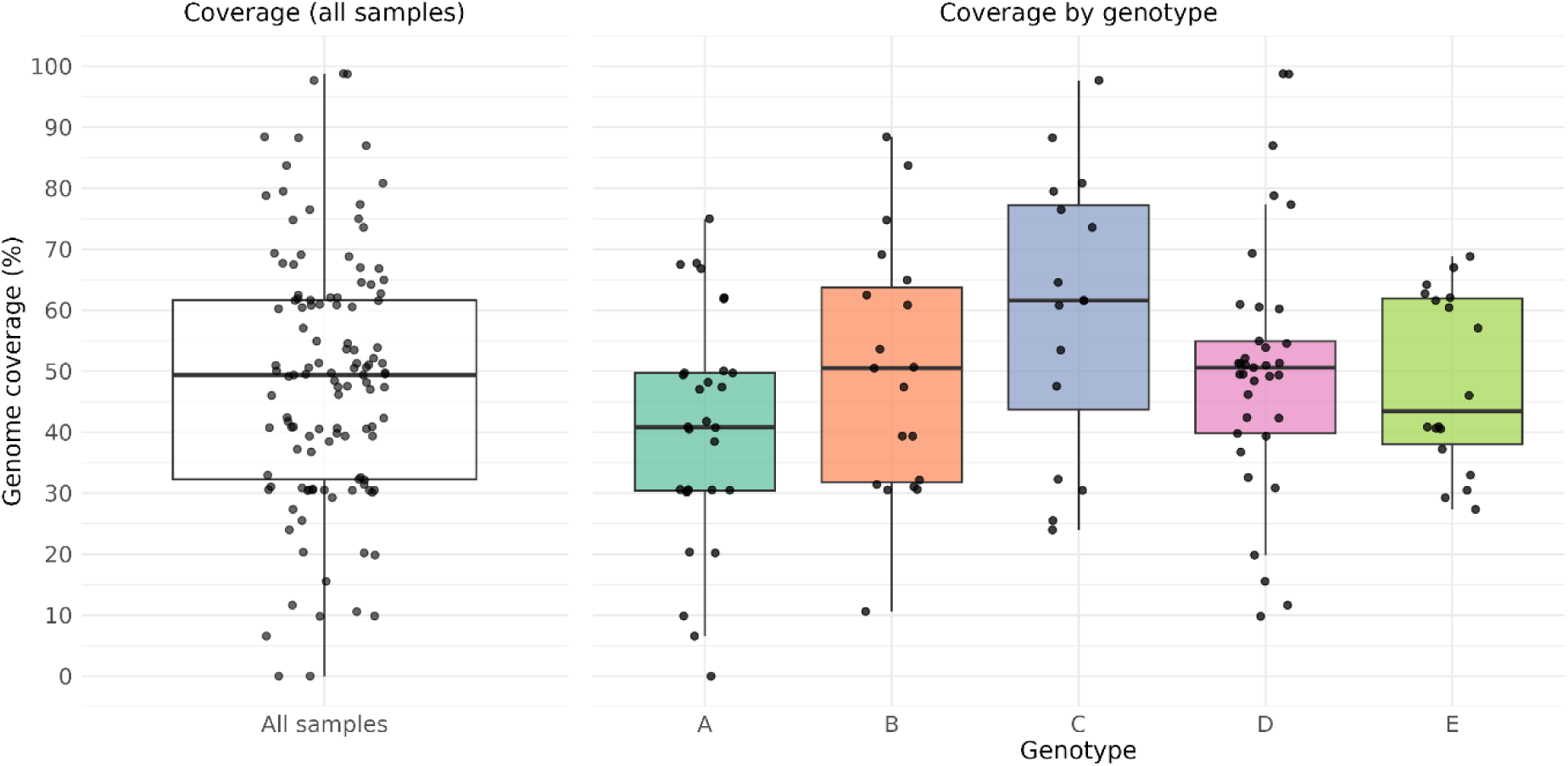
Genome coverage of all samples. Left panel shows the genome coverage in percentage of all samples. The right panel shows the genome coverage in percentage according to genotype. The black dots show the percentage of each individual sample. The median coverage is shown by the thick black line, the boxes show the variation in coverage from the 25^th^ to the 75^th^ percentile, and the whiskers extend to the most extreme points that are within 1.5 times the interquartile range.

### Amplicon performance

We then evaluated the amplification performance of each amplicon (Figure 4A). There was substantial variation across the tiled amplicon scheme, with overall two distinct regions of the genome showing differential amplification success. Amplicons 1-5 generally achieved high sequencing depths, with amplicons 2, 3 and 5 consistently reaching median depths above 100x (note that the numbers in Figure 4A are normalized down to 100X). However, amplicons 1 and 4 displayed considerable variability, with median depths of approximately 40x and 25x respectively, and individual samples ranging from near zero to over 100x. In contrast, amplicons 6, 8, 9, and 10 showed poor amplification, with median depths below 20x and most samples failing to reach the consensus calling threshold. Amplicon 7 exhibited intermediate performance with a median depth of approximately 15x and greater variability than the other poorly performing amplicons.

**Figure 4:**
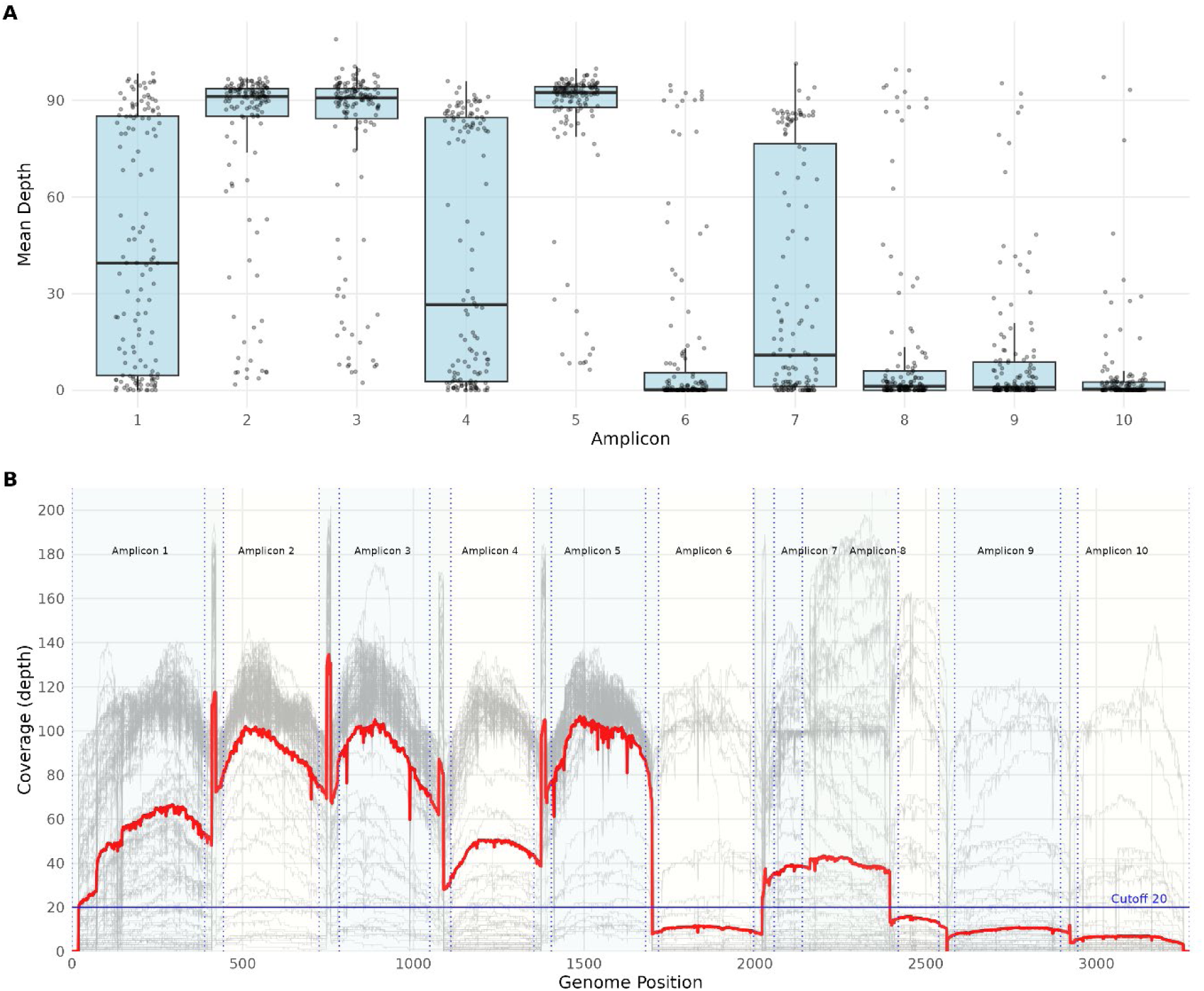
Amplicon performance and genome coverage of all samples. A) Top panel shows the average read depths per sample per amplicon (black dots) with the numbers summarized in the boxplots as in Figure 3. B) Bottom panel shows the read mapping depth per position in the reference genome for all samples (faint grey lines) and the mean depth across all samples as red line. The vertical dashed lines and shaded areas mark the regions amplified by each amplicon. The horizontal blue line at read depth 20 marks the cutoff for calling a consensus nucleotide.

These amplicon-specific differences are reflected in the genome-wide coverage profile (Figure 4B). The substantial overlap between amplicons 7 and 8 contributes to the higher and more variable coverage observed in this region despite the generally poor performance of individual amplicons in the second half of the genome. Notably, regions covered by amplicons 6 and 8–10 frequently fell below the 20x depth threshold required for consensus base calling, which explains the approximately 50% median genome coverage observed across samples.

### Sensitivity and specificity

We examined the relationship between Ct value and genome coverage (Figure 5A). There was a moderate negative correlation between Ct value and genome coverage (r = -0.59, p < 0.001), with Ct values explaining approximately 33% of the variance in coverage (R² = 0.34). Samples with low Ct values (high viral load) generally achieved higher genome coverage, with most samples below Ct 20 reaching 50–80% coverage. In contrast, samples with Ct values above 25 showed highly variable coverage ranging from 0% to 60%, with many falling below 50%.

**Figure 5:**
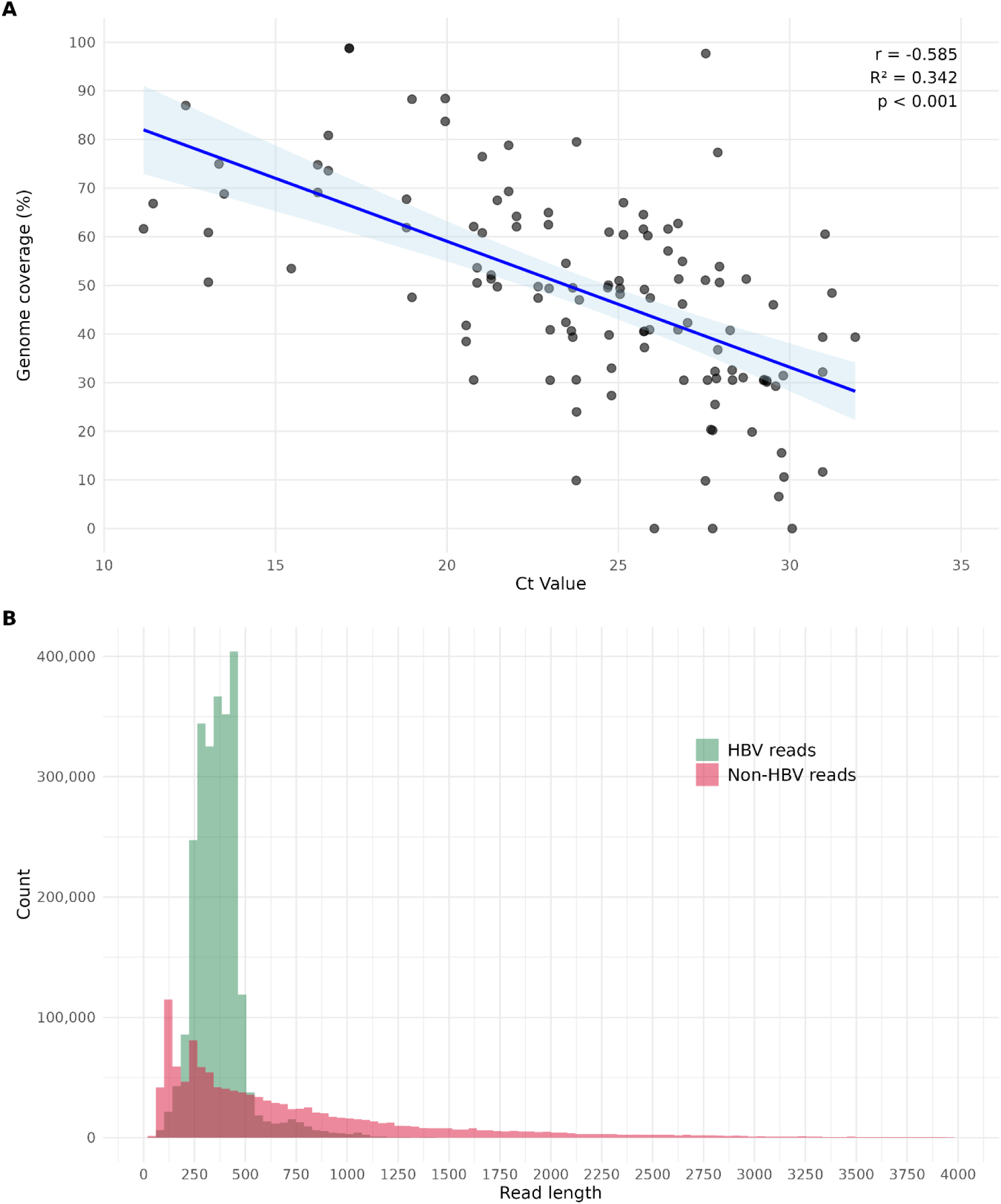
Sensitivity and specificity analysis. A) Scatter plot of genome coverage versus Ct values for all samples, with linear regression line and 95% confidence interval. Pearson correlation coefficient (r), coefficient of determination (R^2^), and p-value are shown. B) Histogram of read length distribution across all samples. Reads classified as HBV by Kraken2 are shown in green; unclassified reads are shown in red.

A similar dependency was observed also for the individual amplicons (Supplementary Figure 1). Across all amplicons, normalized read depth decreased with increasing Ct value, consistent with progressive amplicon dropout at lower template concentrations. Although, the best performing amplicons still amplified well for Ct values above 30, overall there was a trend that the read depths were reduced with higher Ct. However, this effect was highly amplicon-specific. Amplicons 3 and 5 consistently achieved high read depth across a wide Ct range and remained detectable even at Ct values above 30, whereas the poorly performing amplicons, such as amplicons 9 and 10, showed low read depth across nearly all amounts of viral DNA.

To assess the specificity of the primer scheme, we analyzed the read length distribution and taxonomic classification of all sequenced reads (Figure 5B). The vast majority of reads were classified as HBV by Kraken2, demonstrating high specificity of the PCR amplification overall. Non-HBV reads represented only a small fraction of the total. The HBV-classified reads showed a length distribution centered around the expected amplicon sizes, whereas non-HBV reads exhibited a broader length distribution extending to much longer fragments. This pattern suggests that off-target reads may originate from non-specific primer binding. Notably, the HBV read length distribution showed a slight skew toward shorter fragments, which reflects the fragmentation during library preparation.

To further characterize the non-HBV fraction, all samples were additionally screened for human reads by mapping against the human reference genome. Human reads were detected at low levels in most samples (median: 1.8%), though a subset of samples showed markedly elevated human read fractions (Supplementary Table S1 and Supplementary Figure 2). These samples were predominantly characterized by high Ct values, very low total read counts, and poor genome coverage, consistent with failed or poorly performing PCRs rather than true human contamination or non-specific binding. In such samples, the near-absence of amplified HBV DNA means that human background DNA, inevitably present at trace levels from the clinical material, constitutes a disproportionately large share of the sequencing output.

### Primer mismatch analysis

Analysis of primer mismatches across reference sequences from genotypes A–E showed in general low levels of mismatch to the different genotypes, with most primers having on average fewer than one mismatch across the full primer sequence (Table 3). Although some primers showed higher levels of mismatch. When averaged across genotypes, the highest mismatch counts were observed for HBV_4_LEFT, followed by HBV_3_RIGHT, HBV_8_LEFT, and HBV_9_RIGHT. HBV_2_RIGHT, HBV_6_RIGHT, HBV_7_LEFT, and HBV_10_LEFT, also showed slightly elevated mismatch levels. In contrast, most primers displayed fewer than one mismatch, including both primers in amplicon one, and several in the central region of the genome. Importantly, mismatch levels at the critical 3′ terminal five nucleotides were uniformly low across all primers, indicating that the observed mismatches were distributed mainly outside the priming terminus. No consistent genotype-specific pattern was detected.

**Table 3:** Primer mismatch analysis. The mean number of mismatches against the different reference sequences belonging to each genotype is shown (columns A-E), with the mean number of mismatches across all genotypes shown in the “Total” column. The number of mismatches are shown across the entire primer sequence (“Full primer”) and for the last 5 nucleotides (“3’ end”). Numbers are colored from white (zero mismatches) to dark grey (highest number of mismatches) for visualization purposes.

| Primer | Full primer |  |  |  |  |  | 3’ end |  |  |  |  |  |
| --- | --- | --- | --- | --- | --- | --- | --- | --- | --- | --- | --- | --- |
|  | Genotype |  |  |  |  |  | Genotype |  |  |  |  |  |
|  | A | B | C | D | E | Total | A | B | C | D | E | Total |
| HBV_1_LEFT | 0 | 0.2 | 0.2 | 1 | 0 | 0.3 | 0 | 0 | 0 | 0 | 0 | 0 |
| HBV_1_RIGHT | 1 | 1.3 | 0.1 | 0.8 | 0 | 0.6 | 0 | 0 | 0 | 0 | 0 | 0 |
| HBV_2_LEFT | 0 | 0 | 1.1 | 0 | 0 | 0.2 | 0 | 0 | 0 | 0 | 0 | 0 |
| HBV_2_RIGHT | 1 | 1.3 | 1.3 | 1 | 1 | 1.1 | 0 | 0 | 0 | 0 | 0 | 0 |
| HBV_3_LEFT | 1 | 1 | 0 | 0 | 0 | 0.4 | 0 | 0 | 0 | 0 | 0 | 0 |
| HBV_3_RIGHT | 0 | 1.3 | 0.3 | 1.5 | 0 | 0.6 | 0 | 0 | 0 | 0 | 0 | 0 |
| HBV_4_LEFT | 2.1 | 0.7 | 2.5 | 1.8 | 3 | 2 | 0 | 0 | 0 | 0 | 0 | 0 |
| HBV_4_RIGHT | 0.9 | 0.2 | 0.8 | 0.3 | 0 | 0.4 | 0 | 0 | 0 | 0 | 0 | 0 |
| HBV_5_LEFT | 0 | 0 | 0.2 | 0 | 0 | 0 | 0 | 0 | 0.1 | 0 | 0 | 0 |
| HBV_5_RIGHT | 0.1 | 0.2 | 0.1 | 0 | 0 | 0.1 | 0.1 | 0 | 0 | 0 | 0 | 0 |
| HBV_6_LEFT | 1 | 0 | 0.1 | 0 | 0 | 0.2 | 1.0 | 0 | 0 | 0 | 0 | 0.2 |
| HBV_6_RIGHT | 0.5 | 1.2 | 1.1 | 0.3 | 0 | 0.6 | 0 | 0 | 0 | 0 | 0 | 0 |
| HBV_7_LEFT | 1 | 0.3 | 0.4 | 1.5 | 1 | 0.9 | 0 | 0 | 0 | 0 | 0 | 0 |
| HBV_7_RIGHT | 0 | 0 | 0 | 0.6 | 0 | 0.1 | 0 | 0 | 0 | 0.3 | 0 | 0.1 |
| HBV_8_LEFT | 0.8 | 1.2 | 1.2 | 0.6 | 2 | 1.1 | 0 | 0 | 0 | 0.1 | 0 | 0 |
| HBV_8_RIGHT | 1 | 1.3 | 0.2 | 1.3 | 2 | 1.2 | 0 | 0 | 0 | 0 | 0 | 0 |
| HBV_9_LEFT | 0.1 | 0.5 | 1.1 | 0.3 | 1 | 0.6 | 0 | 0 | 0 | 0 | 1.0 | 0.2 |
| HBV_9_RIGHT | 0.5 | 0.7 | 0 | 2.8 | 3 | 1.4 | 0 | 0 | 0 | 0 | 0 | 0 |
| HBV_10_LEFT | 1.3 | 0.8 | 0.2 | 0 | 1 | 0.7 | 0 | 0 | 0.1 | 0 | 0 | 0 |
| HBV_10_RIGHT | 0.5 | 0.5 | 0 | 0.1 | 0 | 0.2 | 0.1 | 0.2 | 0 | 0 | 0 | 0.1 |

### Consensus depth stringency

We also evaluated the potential impact of lowering the minimum depth cutoff from 20x to 10x for consensus base calling. Reducing the threshold increased mean genome coverage from 48.9% to 54.3%, an improvement of 5.5 percentage points. On average, this added 179 bp to consensus sequences (median: 146 bp, range: 0–768 bp), with 56% of samples gaining more than 100 bp. Comparison with Sanger sequences covering roughly amplicon 1-3 showed that the lower cutoff introduced slightly more mismatches (mean 2.92 ± 6.85 SD vs 1.97 ± 4.97 SD SNPs), representing on average only one additional mismatch per sample. However, only 5.4% of all amplicons across all samples had depths between 10–20x, the range where the lower cutoff would include them in the consensus. The underperforming amplicons (6, 8, 9, and 10) predominantly had depths below 10x, indicating that the lower cutoff would not substantially rescue these regions.

## Discussion

In this study we evaluated the feasibility of adapting the (Ringlander et al. 2022) primer scheme, originally developed for singleplex Ion Torrent sequencing, to a multiplexed tiling PCR strategy compatible with Oxford Nanopore sequencing. Our findings demonstrate that while the primer set can be made to function under multiplexed conditions, independent of combining overlapping amplicons in a single reaction or not, performance is limited by recurrent underamplification of specific genomic regions. Failures were concentrated in a reproducible set of genomic intervals and were not clearly linked to genotype or to the choice of primer pooling strategy. The specificity of the amplification reaction was high overall, with the vast majority of reads centered around expected amplicon sizes and classified as HBV, confirming that removal of Ion Torrent adapters did not introduce major off-target amplification. Likewise, human-derived reads were present at low levels in most samples. Only in samples with high Ct values and very low total read counts did we observe substantial levels of human reads. However, these represented failed or poorly performing PCRs.

Across the samples analyzed, genome recovery was modest, with a median of approximately half of the HBV genome represented, although with substantial variation ranging from >90% coverage to complete failure. A pronounced polarity in amplicon performance was consistently observed regardless of genotype, Ct-value or primer pooling strategy. Amplicons 1–5 amplified reasonably well, while the remaining amplicons (perhaps except for amplicon 7) often failed to reach consensus-level depth. Even when the consensus depth cutoff was reduced from 20x to 10x. This asymmetric behavior explains the characteristic coverage plateau around 50% observed across samples.

The decline in genome recovery with increasing Ct value reflects progressive dropout of individual amplicons rather than a uniform reduction in coverage across the genome. As Ct values increased, robust amplicons continued to amplify across a wide Ct range, while weaker amplicons dropped out early. The consistent amplification of amplicons 3 and 5 across nearly the full Ct range explains why partial genomes were frequently recovered even at low levels of viral DNA.

The observed amplification pattern cannot be explained by genotype-specific effects. Across samples, multiple genotypes showed similar coverage profiles and the same subsets of amplicons repeatedly succeeded or failed. The polarity of amplification performance, together with the fact that all amplicons occasionally reached high depths, points to limitations in primer design and primer-template interaction.

Primer mismatch analysis against the different genotypes did not reveal a consistent relationship between number of mismatches and amplification performance. Overall, mismatch counts were generally low across genotypes and showed no clear threshold beyond which amplification systematically failed. While some correspondence was observed for individual regions, for example the primers for amplicons 4, 8, and 9 showed elevated mean mismatch counts and these amplicons were among those with the most unstable or frequently reduced coverage. This relationship was not generalizable and several poorly performing amplicons, such as amplicon 6 and 10, had few mismatches across genotypes.

Thus, while primer–template mismatches likely contribute to reduced amplification efficiency in specific regions, mismatch burden alone does not predict amplicon performance. Nevertheless, there is room for optimalization of the primer sequences. In particular, improving consistently weak amplicons, rather than increasing the sequencing depth or relaxing the coverage thresholds, would be expected to greatly improve the sensitivity of the assay.

The most consistently recovered genome region comprised the first ∼800 nt of the reference, corresponding to amplicons 1–3. This region overlaps the polymerase reverse-transcriptase domain and the major surface antigen and is commonly used for HBV genotyping and clinical resistance testing. Thus, even when full-genome recovery is incomplete, the parts of the genome most relevant to surveillance and resistance monitoring are usually obtained.

Importantly, multiplexing itself did not introduce major additional bias. Genome coverage and normalized amplicon depths were largely correlated between primer pooling strategies. Nevertheless, while avoiding amplifying overlapping amplicons in the same reaction yielded marginally higher coverage, neither approach overcame the fundamental coverage instability imposed by weak amplicons. Given this, in high-throughput laboratory workflows, such as in surveillance programs, pooling all primers into a single reaction may be an acceptable trade-off. It substantially reduces the workload and opportunities for error while yielding similar results as separating the non-overlapping amplicons.

Compared to more recent HBV whole-genome amplification strategies, the limitations of the Ringlander et al. (2022) scheme become particularly clear. Newer approaches, including the scheme described by Lumley et al. (2025), achieve substantially higher and more uniform genome recovery through redesigned tiling strategies specifically optimized for multiplex PCR and long-read sequencing. While Lumley at al. (2025) use fewer and longer amplicons than the Ringlander et al. primers, which thereby should be more sensitive to for example degraded template DNA, an important difference to their primer scheme is that they designed multiple overlapping or degraded primers per binding site. This in effect makes the primer scheme cover more of the potential HBV genomic diversity.

Our study, together with other recent tiling amplicon amplifications of HBV, shows that multiplex tiling PCR amplification of entire HBV genomes is feasible and can be implemented as a cost-effective and rapid method in HBV genotyping and surveillance.

## Supplementary material

**Supplementary Table S1.** Table containing for each analyzed sample the name of the Nanopore sequencing run (“Run”), barcode (“Barcode”), sample ID (“Sample_ID”), primer pooling strategy (“Pooling_strategy”), Ct value (“Ct_value”), predicted genotype of the Sanger sequenced fragment (“Sanger_genotype”), number of raw Nanopore reads (“Raw_read_count”), number of reads removed when mapping to the human genome (“Human_reads_removed”), percentage of human reads out of the total reads (“Human_pct”), number of mapped reads (“Mapped_read_count”), percentage of mapped reads out of the total reads (“Mapped_pct”), number of HBV classified reads (“Kraken_classified_reads”), and the percent of the reference genome covered by the consensus sequence (“Coverage_breadth”).

---

**Supplementary Figure 1.**
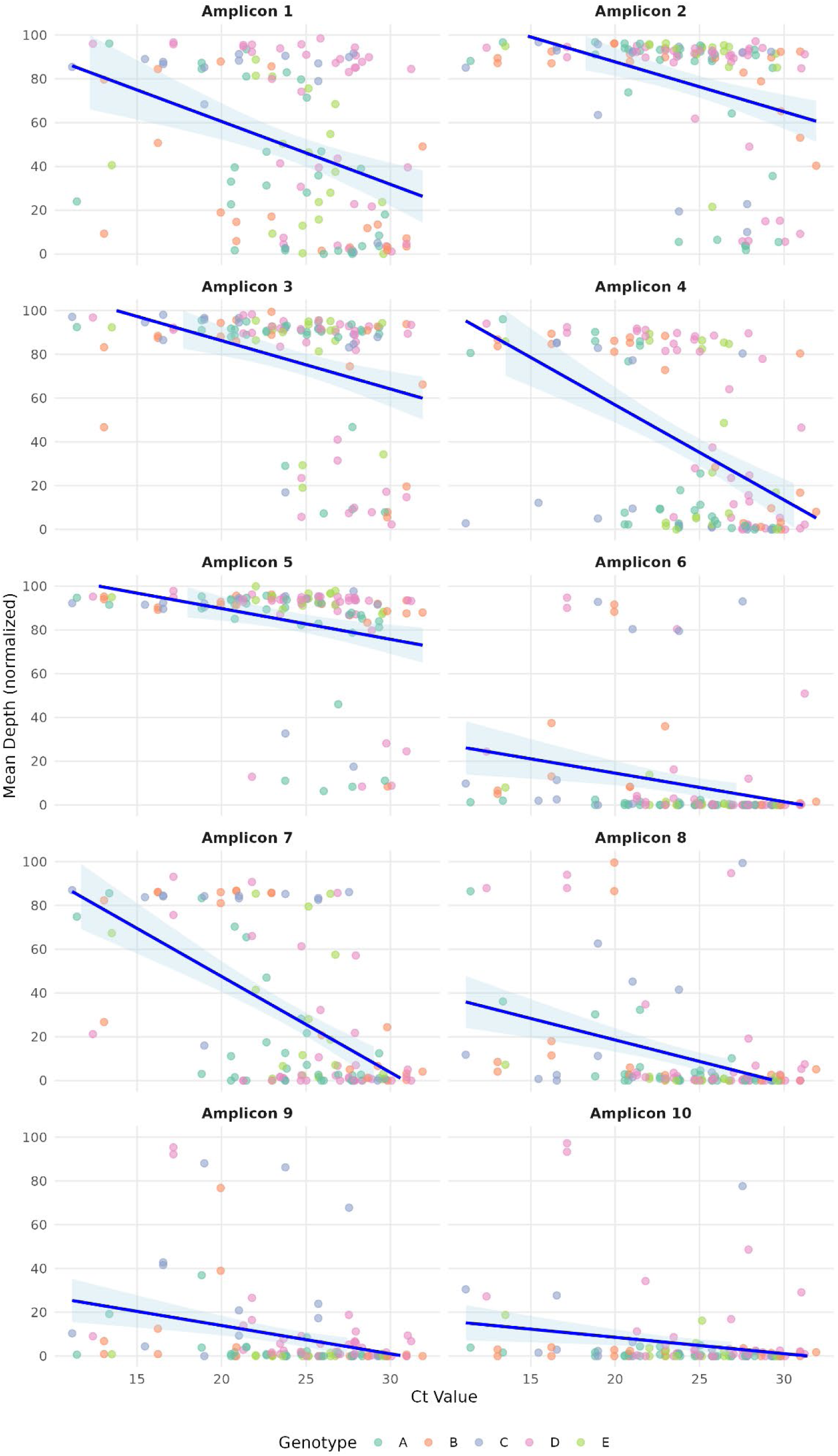
A) Scatter plot of normalized read count (from the ARTIC pipeline) per amplicon (y-axis) versus Ct values (x-axis) for all samples, with linear regression line and 95% confidence interval. Pearson correlation coefficient (r), coefficient of determination (R^2^), and p-value are shown. Samples are colored according to genotype.

**Supplementary Figure 2.**
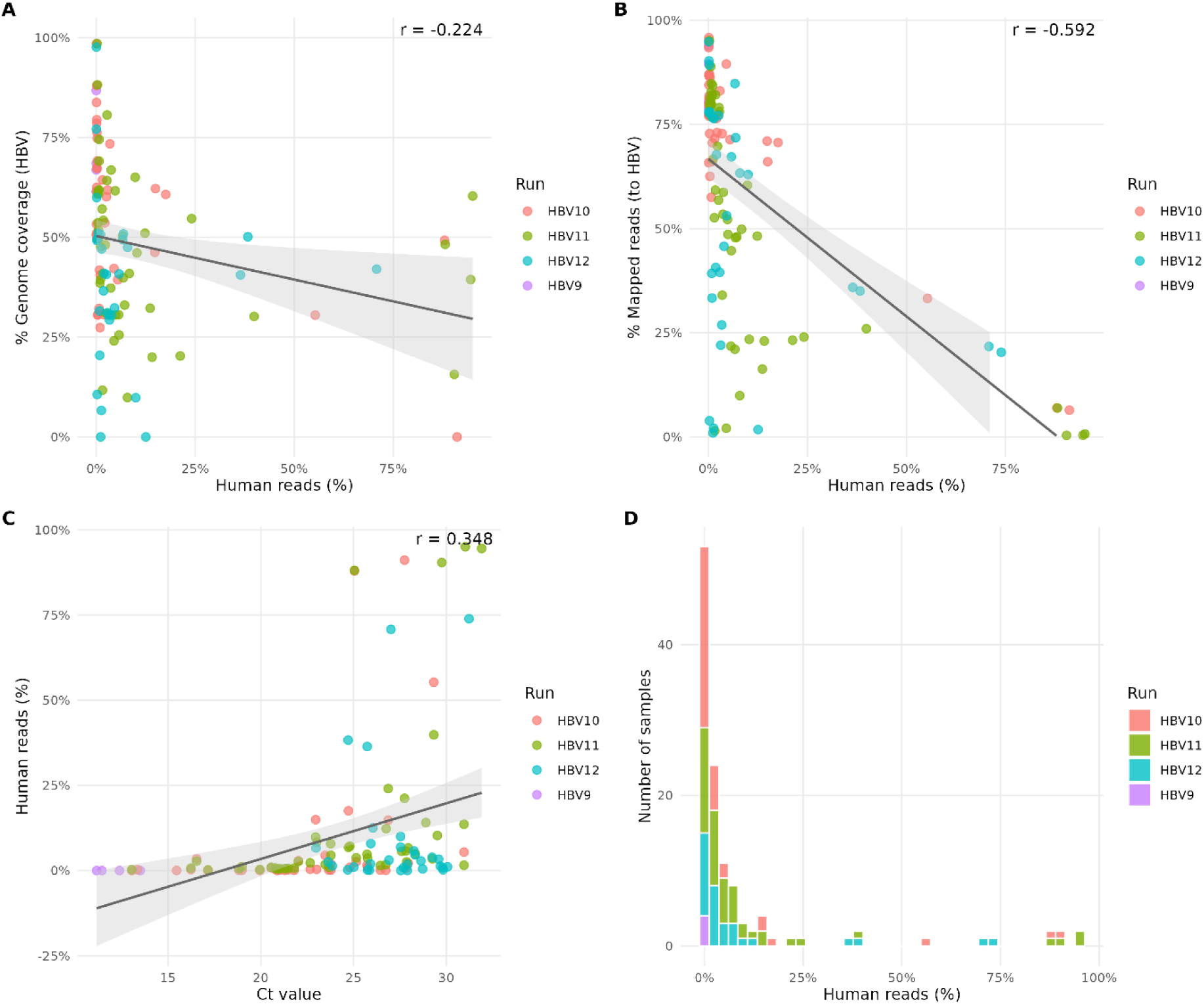
(A) Percentage of human reads (identified by read mapping against the human reference genome) plotted against HBV genome coverage breadth (%) for each sample. (B) Percentage of mapped human reads per sample versus the percentage of reads mapping to the HBV reference genome. (C) Relationship between sample Ct value and percentage of mapped human reads. (D) Distribution of human read percentages across all samples, stratified by Nanopore sequencing run. In all panels, each point represents one sample, coloured by sequencing run. Regression lines with 95% confidence intervals are shown in grey. Pearson correlation coefficients (r) are indicated in each panel.

